# RAAS blockade, kidney disease, and expression of *ACE2*, the entry receptor for SARS-CoV-2, in kidney epithelial and endothelial cells

**DOI:** 10.1101/2020.06.23.167098

**Authors:** Ayshwarya Subramanian, Katherine A Vernon, Michal Slyper, Julia Waldman, Malte D Luecken, Kirk Gosik, Dan Dubinsky, Michael S Cuoco, Keith Keller, Jason Purnell, Lan Nguyen, Danielle Dionne, Orit Rozenblatt-Rosen, Astrid Weins, Human Cell Atlas Lung Biological Network, Aviv Regev, Anna Greka

## Abstract

SARS-CoV-2, the coronavirus that causes COVID-19, binds to angiotensin-converting enzyme 2 (ACE2) on human cells. Beyond the lung, COVID-19 impacts diverse tissues including the kidney. ACE2 is a key member of the Renin-Angiotensin-Aldosterone System (RAAS) which regulates blood pressure, largely through its effects on the kidney. RAAS blockers such as ACE inhibitors (ACEi) and Angiotensin Receptor Blockers (ARBs) are widely used therapies for hypertension, cardiovascular and chronic kidney diseases, and therefore, there is intense interest in their effect on ACE2 expression and its implications for SARS-CoV-2 pathogenicity. Here, we analyzed single-cell and single-nucleus RNA-seq of human kidney to interrogate the association of ACEi/ARB use with *ACE2* expression in specific cell types. First, we performed an integrated analysis aggregating 176,421 cells across 49 donors, 8 studies and 8 centers, and adjusting for sex, age, donor and center effects, to assess the relationship of *ACE2* with age and sex at baseline. We observed a statistically significant increase in *ACE2* expression in tubular epithelial cells of the thin loop of Henle (tLoH) in males relative to females at younger ages, the trend reversing, and losing significance with older ages. *ACE2* expression in tLoH increases with age in females, with an opposite, weak effect in males. In an independent cohort, we detected a statistically significant increase in *ACE2* expression with ACEi/ARB use in epithelial cells of the proximal tubule and thick ascending limb, and endothelial cells, but the association was confounded in this small cohort by the underlying disease. Our study illuminates the dynamics of *ACE2* expression in specific kidney cells, with implications for SARS-CoV-2 entry and pathogenicity.

## Introduction

Severe acute respiratory syndrome coronavirus 2 (SARS-CoV-2), the coronavirus responsible for the Coronavirus Disease 2019 (COVID-19) pandemic, enters human cells through the binding of its spike (S) protein to the host cell surface angiotensin-converting enzyme 2 (ACE2) (1, 2). Subsequent cleavage by one of several accessory host proteases, including transmembrane protease serine 2 (TMPRSS2) and cathepsin L (CTSL), is required for successful viral entry and replication (3). While the nasal, lung and gut epithelia are primary targets of SARS-CoV-2 infection, complications affecting multiple organ systems suggest the virus impacts other tissues, and studies have described RNA expression of *ACE2* in diverse tissues including the gut, heart, liver, kidney, olfactory epithelium, brain and vasculature (4, 5). To date, little is known about the cell-type specific expression patterns of *ACE2* in human kidney in a large enough cohort to allow us to account for factors such as age and sex that appear to be critical for susceptibility to SARS-CoV-2 and severity of disease.

ACE2 is a key component of a more recently recognized counter-regulatory arm of the Renin-Angiotensin-Aldosterone system (RAAS (6)), critical for regulating blood pressure. *ACE2* is located on the X chromosome, and RAAS activity has been reported to exhibit sex-specific differences (7, 8). ACE2 converts angiotensin II (Ang II) to the vasodilatory peptide Ang-(1-7), angiotensin I to Ang-(1-9) (also cleaved to Ang-(1-7) by angiotensin converting enzyme (ACE) or neprilysin), and angiotensin A to alamandine (9), a peptide mediating similar actions as Ang-(1-7). Together, this non-classical arm of the RAAS counteracts the vasoconstrictive, proliferative and inflammatory effects of the classical RAAS, mediated by Ang II (generated from angiotensin I by the action of ACE). ACE2 therefore acts as a critical regulator of the Ang II versus Ang-(1-7) balance to prevent deleterious high blood pressure and tissue inflammation, especially in the kidney, where ACE2 has been previously detected at the brush borders of tubular epithelial cells and at lower levels within glomeruli (10).

The RAAS is frequently targeted in the treatment of hypertension and chronic kidney disease (CKD). Common drug modulators of RAAS include ACE inhibitors (ACEi) and Angiotensin Receptor Blockers (ARBs). ACEi block the conversion of Angiotensin I to Ang II by ACE, while ARBs block the binding of Ang II to Ang II receptor type 1 (AT1R), and inhibit its vasoconstrictive effects. The association between ACE2 expression and activity with ACEi/ARB use is not clearly defined (11); results largely depend on the model system (rodents, humans), the specific drug tested, and the tissue of investigation (12–15).

COVID-19 is associated with greater disease severity and mortality in older individuals, in men, and in those with comorbidities, including diabetes mellitus and CKD (16, 17). Since ACE2 acts as the receptor for SARS-CoV-2, there has been widespread debate (18, 19) if altered ACE2 expression and RAAS regulation in patients taking ACEi/ARBs affects the pathogenicity of SARS-CoV-2 infection. Two recent large population-based, case-control studies found no significant association between the use of RAAS blockers and risk or severity of COVID-19, regardless of sex or age (20, 21). However, disease severity was narrowly defined in these studies based on clinical manifestations affecting the lung. Thus, there remains an urgent need for a deeper understanding of the dynamics and relationship between ACE2 expression and ACEi/ARB use at the level of individual cells, and adjusted for age, sex and other confounders in other organ systems, especially the kidney.

To investigate whether ACE2 expression in the kidney, a key RAAS tissue, may be influenced by ACEi/ARB use, we examined transcriptomic data that we had collected at the level of individual kidney cells to answer two key questions: (1) what are the effects of age and sex on *ACE2* expression at the single cell level in the human kidney? (2) Is there a link between the use of ACEi/ARB and *ACE2* expression in the kidney?

## Results and Discussion

### A cross-study integrated-analysis of kidney cells associates *ACE2* expression with age and sex in the thin loop of Henle

We investigated *ACE2* expression in the kidneys of individuals on RAAS blockers, by examining single-cell and single-nucleus RNA-seq (sc/snRNA-seq) data, accounting for age and sex as covariates in the statistical analysis. We first generated a reference atlas of kidney with no known pathology (in the tissue sampled) for the association of *ACE2* expression in individual kidney cell types with sex and age, aggregating 8 independent scRNA-seq or snRNA-seq studies from 8 independent centers (**Dataset S1**) for a total of 176,421 cells profiled from 49 donors (20 females, 29 males), and whose median age was 57 with an IQR of 14. We co-embedded the combined “integrated analysis” dataset adjusting for donor batch effect, and then performed graph-based clustering to derive 24 broad cell subsets (**Methods, Figure S1A**), which we annotated *post hoc*, including podocytes, proximal, distal and collecting duct tubular epithelial cells, immune and stromal cells (**Methods, Figure S1B**), of which 15 subsets had cells from at least 40 donors. No cell subset segregated by sex.

Overall, the proportion of cells expressing *ACE2* (*ACE2*^+^) varied by cell subset from 0% to 9.87% (**Figure S1B**). To account for technical variability, we examined the *ACE2^+^* proportion within each donor per cell subset. The thin loop of Henle (tLoH) had the highest mean percentage (13.7%) of *ACE2^+^* cells per donor, followed by proximal convoluted tubular cells (PCT, 7%), parietal epithelial cells (PEC, 4.6%) and vascular smooth muscle cells (vSMC, 3.5%) (**Figure 1A**).

**Figure 1.**
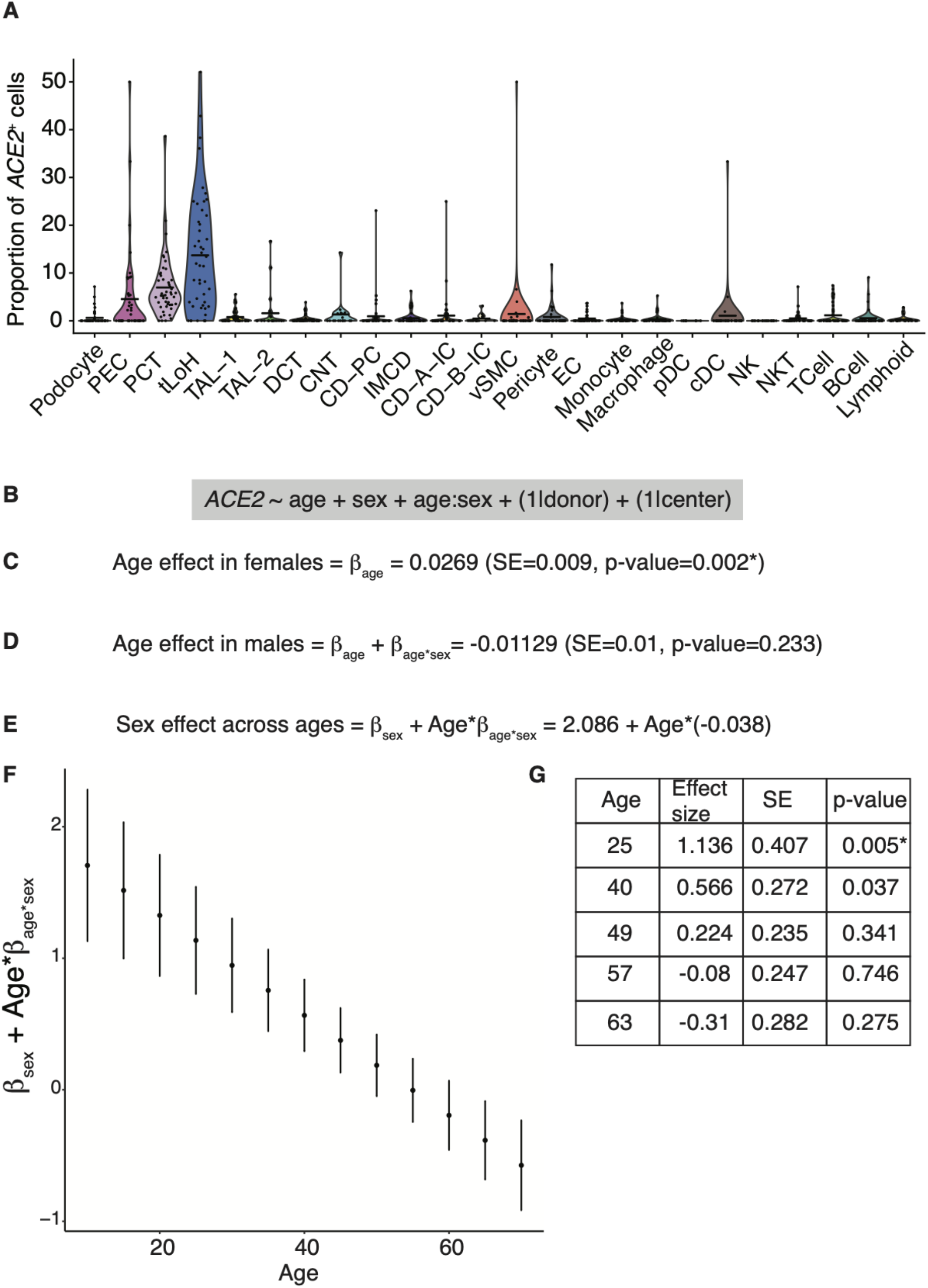
Integrated analysis of 8 reference kidney single-cell datasets shows *ACE2* expression is associated with age, sex and their interaction in the thin loop of Henle tubular epithelial cells of the kidney. (**A**) *ACE2*^+^ cells across kidney cell subsets. Distribution of proportion of cells expressing *ACE2* (count of at least 1) in each individual (dot, *y* axis) in each cell subset (*x* axis). Two outliers were removed, each having only 1 cell of the cell class. Cell proportions were computed as the ratio of the number of cells in a certain cell type expressing *ACE2* divided by the total number of cells in the cell type. (**B**) Statistical model fitted to the data to assess sex, and age associations with gene expression of *ACE2*. (**C**) Age effect in females in the tLoH (**D**) Age effect in males in the tLoH (**E**) Sex effect in tLoH, varies with age (**F**) Scatter plot of sex effect (y-axis) with age (x-axis) from 10y to 70y in increments of 5y. Standard error bars are shown. (**G**) Sex effect computation for younger ages (25, 40 years), first quartile (49y), median age (57y) and second quartile (63y). Asterix (*) indicates statistical significance at FDR of 5%.

To associate *ACE2* expression with donor age and sex, we fit a mixed effect Poisson regression model to cell subsets with at least 40 *ACE2*^+^ cells across donors, with age, sex and their interaction as covariates, and nested random effects for donors and center (**Methods, Figure 1B**). We found a statistically significant association (FDR-adjusted p value < 0.05) between age, sex and *ACE2* expression in the tLoH cell subset (**Figure S2A, Dataset S2**). *ACE2* expression trended upwards with age in females (**Figure 1C**). In males, *ACE2* expression trended downwards with age, and was not statistically significant (**Figure 1D, Methods**). Overall, the sex effect was statistically significant at younger ages, where males had higher expression relative to females. With older ages, the gap narrows, eventually reversing direction and is no longer significant (**Figure 1E, 1F, 1G**). We performed cross-validation to check for robustness of the results to dataset origin, using our recently developed strategy (5) (**Methods, Dataset S2**). The direction of association for age, sex and their interaction for tLoH was preserved and consistent in the cross-validation analysis, except that excluding the “Sanger’’ dataset removed their statistical significance (**Dataset S2**). The Sanger dataset is the largest by number of donors (13 donors; 7 females, 6 males), widest age range (2y to 72y) and second largest in number of cells (40,198 cells). To capture subtler effects, we fit a simpler fixed effects Poisson regression model with age, sex, their interaction and dataset as covariates, alongside a pseudo-bulk analysis to validate that single-cell effects are consistent on the donor level, and cross validation (5) (**Methods, Dataset S2**). A caveat of the integrated analysis is that aggregating cells to broader cell categories across various batches may lead to loss of within-class cellular heterogeneity. Effect sizes may differ across the subclasses, thus impacting their estimation. In conclusion, at baseline, *ACE2* expression had a statistically significant association with both age in females and sex at younger ages in the tLoH cells. Because of the higher median age in our integrated analysis dataset, larger cohorts with wider age ranges are necessary to validate the results.

### Association of RAAS blockade with *ACE2* expression in the kidney epithelial and endothelial cells confounded by underlying disease

Next, to test the association of renal *ACE2* expression with ACEi/ARB use, we surveyed *ACE2* expression in 32,239 nuclei obtained by droplet-based snRNA-seq of frozen kidney samples from an independent cohort of 11 patients: 9 kidney biopsies with features of various kidney diseases including Lupus nephritis (LN) and IgA nephropathy (IgAN) (Table 1) and 2 cortical samples from a tumor nephrectomy and transplant nephrectomy with rejection and recurrent Focal Segmental Glomerulosclerosis (FSGS). 6 of the 9 biopsied patients were either on Lisinopril (ACEi) or Losartan (ARB). Nuclei were isolated using one of two protocols (**Methods, Table 1**). Graph based clustering and post hoc annotation (**Methods**) identified 14 broad cell classes (**Figure S3A**), including epithelial, immune and stromal cells, of which 8 classes had greater than 5 nuclei expressing *ACE2* (**Figure 2A**). In contrast to the baseline analysis, PCT cells had the highest proportion of *ACE2^+^* cells across this patient cohort, we did not recover tLoH nuclei, and PECs had fewer than 5 *ACE2^+^* nuclei. To account for technical sampling differences, we analyzed *ACE2+* cell proportions for each individual patient (**Figure 2B**).

**Figure 2.**
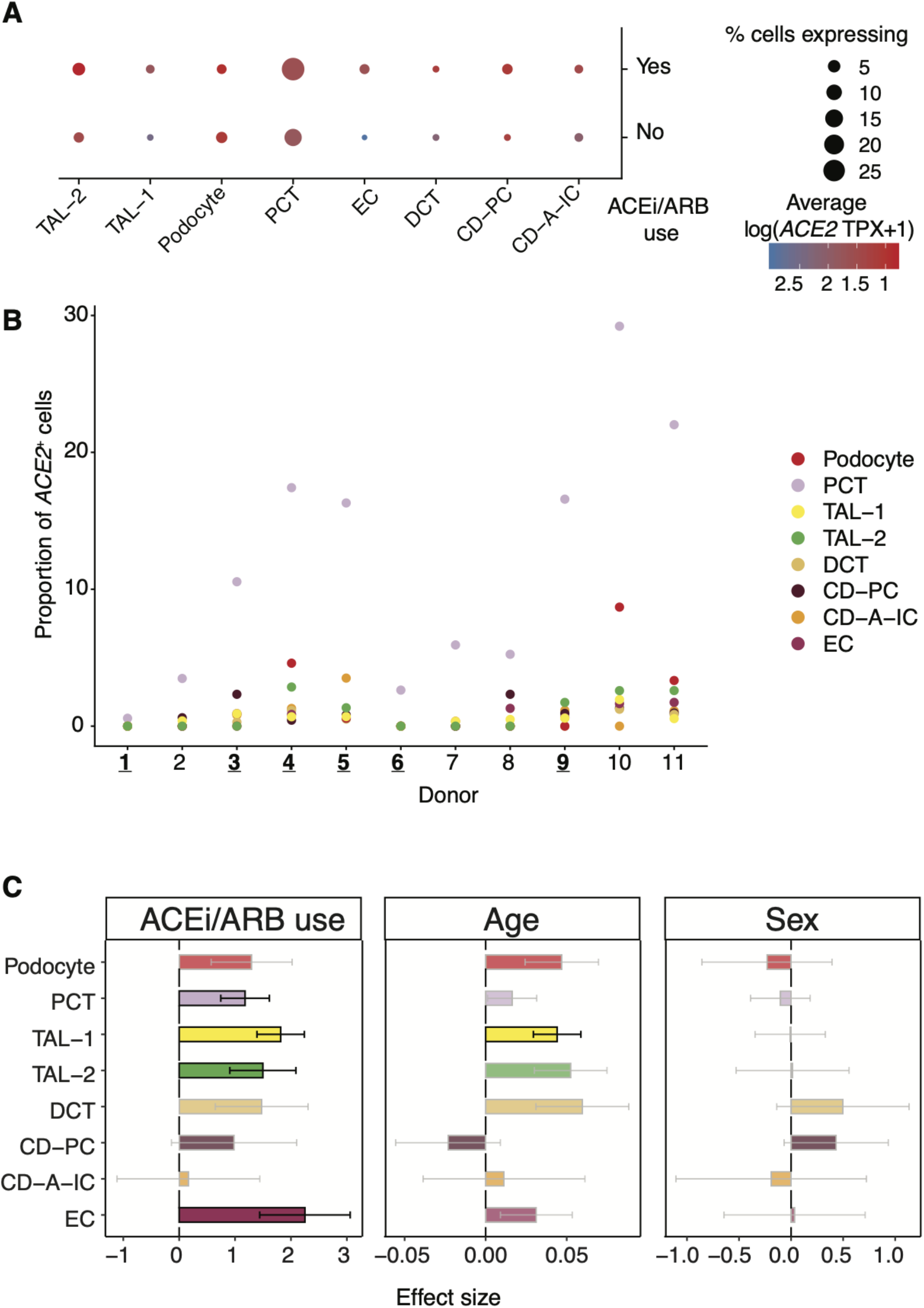
Association of *ACE2* expression with ACEi/ARB use in kidney proximal tubular, thick ascending limb and endothelial cells. (**A**) ACE2 expression across kidney cell subsets in tissue biopsies. Mean *ACE2* expression in *ACE2+* cells (log(TPX+1), dot color) and proportion of expressing cells (dot size) across cell types (columns) with at least 5 *ACE2+* nuclei, stratified by ACEi/ARB use (rows). (**B**) Proximal tubular cells have the highest proportion of ACE2+ cells across patients. Proportion of *ACE2+* cells (y axis) in each cell type (color) in each donor (*x* axis). Donors on RAAS blockade are in bold and underlined (**C**) Association between *ACE2* expression and ACEi/ARB while controlling for age, and sex. The x-axis is the effect size of the association in log-fold change (sex), or slope of log expression with age. Error bars represent standard errors around coefficient estimates. Statistically significant associations (FDR 5%) are outlined in black, others are in gray and colors are shaded lighter.

**Table 1.**
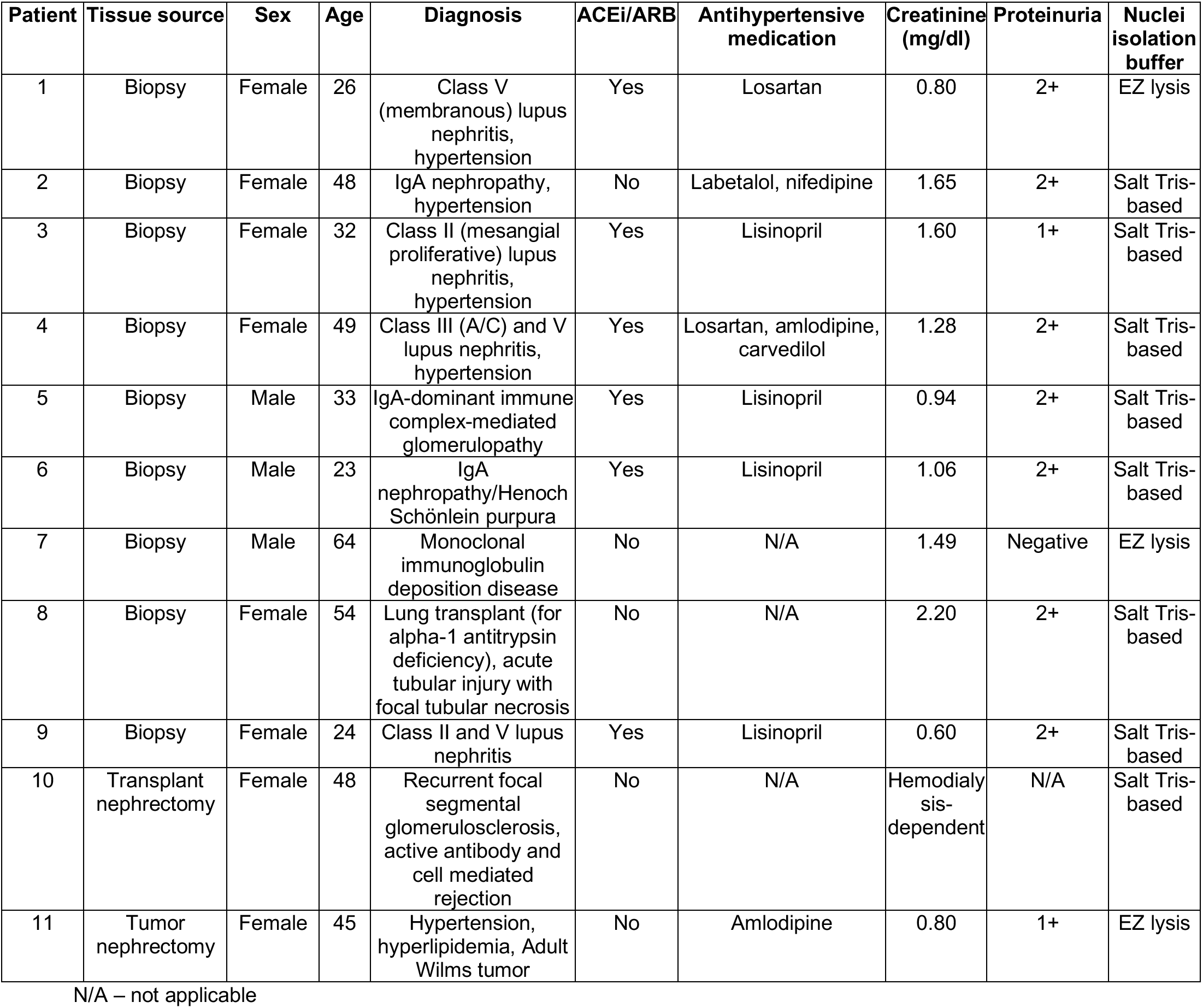
Demographic patient data and nuclei isolation method.

| Patient | Tissue source | Sex | Age | Diagnosis | ACEi/ARB | Antihypertensive medication | Creatinine (mg/dl) | Proteinuria | Nuclei isolation buffer |
| --- | --- | --- | --- | --- | --- | --- | --- | --- | --- |
| 1 | Biopsy | Female | 26 | Class V (membranous) lupus nephritis, hypertension | Yes | Losartan | 0.80 | 2+ | EZ lysis |
| 2 | Biopsy | Female | 48 | IgA nephropathy, hypertension | No | Labetalol, nifedipine | 1.65 | 2+ | Salt Tris-based |
| 3 | Biopsy | Female | 32 | Class II (mesangial proliferative) lupus nephritis, hypertension | Yes | Lisinopril | 1.60 | 1+ | Salt Tris-based |
| 4 | Biopsy | Female | 49 | Class III (A/C) and V lupus nephritis, hypertension | Yes | Losartan, amlodipine, carvedilol | 1.28 | 2+ | Salt Tris-based |
| 5 | Biopsy | Male | 33 | IgA-dominant immune complex-mediated glomerulopathy | Yes | Lisinopril | 0.94 | 2+ | Salt Tris-based |
| 6 | Biopsy | Male | 23 | IgA nephropathy/Henoch Schönlein purpura | Yes | Lisinopril | 1.06 | 2+ | Salt Tris-based |
| 7 | Biopsy | Male | 64 | Monoclonal immunoglobulin deposition disease | No | N/A | 1.49 | Negative | EZ lysis |
| 8 | Biopsy | Female | 54 | Lung transplant (for alpha-1 antitrypsin deficiency), acute tubular injury with focal tubular necrosis | No | N/A | 2.20 | 2+ | Salt Tris-based |
| 9 | Biopsy | Female | 24 | Class II and V lupus nephritis | Yes | Lisinopril | 0.60 | 2+ | Salt Tris-based |
| 10 | Transplant nephrectomy | Female | 48 | Recurrent focal segmental glomerulosclerosis, active antibody and cell mediated rejection | No | N/A | Hemodialysis-dependent | N/A | Salt Tris-based |
| 11 | Tumor nephrectomy | Female | 45 | Hypertension, hyperlipidemia, Adult Wilms tumor | No | Amlodipine | 0.80 | 1+ | EZ lysis |

N/A – not applicable

Because the number of observations per cell type was not large enough to consistently fit a complex model that accounts for both donor variability as a random effect and an interaction between age and sex, we fit a Poisson mixed effects model of *ACE2* gene expression counts in cell classes with greater than 5 *ACE2+* nuclei, with the donor as the random covariate to account for batch effects, while adjusting for age and sex as covariates but not their interaction or center effect as all samples came from the same cohort **(Methods, Dataset S3, Figure 2C)**. In all cases, we sampled cells so that the distributions of the number of Unique Molecular Identifiers (UMIs) were matched between patients on and off ACEi/ARBs to account for sequencing depth differences.

ACEi/ARB use was associated (FDR-adjusted p value < 0.05) with increased *ACE2* expression in select epithelial cells (PCT, Thick Ascending Limb (TAL)) and in endothelial cells (EC). There was also a statistically significant increase in *ACE2* expression with age in the TAL-1 cells. To capture more subtle effects, we also fit a simpler fixed effects model, without accounting for donor effect, but with an interaction term between sex and age, alongside a pseudo-bulk analysis ((5), **Methods**). The direction of the effect and statistical significance for ACEi/ARB use and age were preserved in PCT, TAL and EC. The pseudo-bulk analysis also preserved the direction of effect sizes. We found no new effects with the simpler model (**Dataset S3**). The biopsy cohort differs from the baseline cohort in having a lower median age (45 vs 57), wider age range (IQR of 19.5), and a higher female to male ratio (8:3). Taken together, neither the baseline nor the biopsy cohort analysis found a robust association between *ACE2* expression and sex or age in classical PCT cells. Importantly, the biopsy cohort showed a higher expression of *ACE2* in PCTs, TAL and ECs in patients taking ACEi/ARBs.

A limitation of the study is the small sample size of the biopsy cohort which does not allow us to adequately model the underlying kidney pathology without confounding with RAAS blockade treatment. In particular, as donors had a variety of different kidney diseases, we grouped them into broad disease categories: IgAN (n=3), LN (n=4) and Others (n=4). Because patients in the “Others” category were not treated with ACEi/ARB, the disease and treatment are confounded. Accordingly, when we fit the Poisson model with donor as random effect, and RAAS blockade, age, sex as fixed effects but included an additional fixed covariate for disease category (**Methods**), the association with RAAS no longer held (consistent with confounding), but the age effect in TAL-1 was preserved. In a simpler model with just the donor effect and disease status, LN was associated with *ACE2* expression in PCT, EC and TAL (**Dataset S3**). Our results highlight the need for further investigation with disease stratification and with greater power, to verify if the disease (LN) and RAAS blockade effects counter each other or if there is a true disease effect. In addition to the impact of underlying kidney pathology, other caveats to address in larger studies are ACEi/ARB dose variability and duration of use (although all patients were normotensive at the time of tissue collection), the impact of experimental protocols (freezing, storage and nuclei isolation), and sampling variability in cell composition and depth of sequencing.

Reported manifestations of kidney involvement in COVID-19 patients range from urinary abnormalities and mild functional impairment, to severe acute kidney injury necessitating renal replacement therapy, associated with excess mortality risk (22). In addition to potential effects of ischemia and the cytokine storm associated with the systemic response to SARS-CoV-2 infection, there are reports of a direct viral cytopathic effect mediating kidney tubular injury (23–26). Postmortem examination of kidney tissue from COVID-19 patients revealed evidence of SARS-CoV/SARS-CoV-2 nucleocapsid protein in tubular epithelium and coronavirus particles in the cytoplasm of the proximal, and to a lesser extent distal, tubular epithelium (23, 24). Features suggestive of intracellular virus assembly were also observed (25). Prominent ACE2 staining has also been seen in the proximal tubules of COVID-19 patients, along with focal parietal epithelial cell staining (and occasional weaker podocyte staining) (23). In one report, tubular SARS-CoV-2 was accompanied by tubulointerstitial macrophage infiltration and tubular C5b-9 deposition (24). These data raise the possibility that viral entry into the kidney and any downstream inflammation may be exaggerated when ACE2 expression is upregulated.

In sum, our study provides data at the single-cell level in the kidney, an organ with observed SARS-CoV-2 tropism. Our integrated analysis revealed a statistically significant association between *ACE2* expression, and age and sex in the thin loop of Henle. The varying trends in *ACE2* expression with age and sex in the kidney imply a more complex relationship between gene expression and simple demographic variables like age and sex, which needs further investigation in larger cohorts with wider-ranging ages. The use of ACEi/ARBs was likely confounded with underlying Lupus nephritis in relation to *ACE2* levels in the proximal tubular, thick ascending limb and endothelial cells in the kidney after adjustment for age and sex. Assessing if increased *ACE2* expression is beneficial or harmful in settings of disease or RAAS blockade, requires further mechanistic investigation, in addition to studies in larger patient cohorts. Whether such transcriptional changes also play a role in SARS-CoV-2 tropism for the kidney remains an open question that we may only be able to confidently answer once large clinical cohorts are analyzed at the end of the current pandemic. Further, all our data comes from non-COVID-19 samples, and the effect of virus infection on ACE2 expression is yet to be determined.

## Materials and methods

### Datasets used in the baseline integrated analysis

We included 8 datasets in the integrated analysis for a total of 176,421 cells **(Dataset S1)** post quality control filtering (see below). In addition to an internal unpublished single-cell cohort, we aggregated control kidney single-cell or single-nucleus RNA-seq data from 7 published studies for a total of 49 donors (29 males, 20 females) with a median age of 57 (min 2y, max 72y, IQR=14). Public datasets were obtained at the level of gene expression counts post genome alignment and gene-level quantification as published by each study’s authors (**Dataset S1**). For 2 donors (AMP dataset), only age ranges were available, so mean ages were computed.

### Single nucleus isolation from human kidney tissue

Frozen kidney biopsy specimens or frozen samples of macroscopically normal cortex from tumor nephrectomies (distant from the tumor site), were obtained after appropriate patient (discarded tissue) consent and in accordance with Partners Healthcare IRB and institutional guidelines. One of two protocols (utilizing either nuclei EZ lysis buffer (Sigma-Aldrich, St. Louis, USA) or a salt Tris-based buffer (27, 28)) was used to isolate nuclei from these; in the case of kidney biopsy tissue, the surrounding OCT embedding medium was first removed using PBS. 8,000 or 10,000 single nuclei were loaded into each channel of the Chromium single cell 3’ chip, according to use of the v2 or v3 kit respectively (10x Genomics, Pleasanton, USA).

### Single cell isolation from human kidney tissue

Samples of macroscopically normal cortex were obtained from tumor nephrectomy specimens, distant from the tumor site, after appropriate patient consent and in accordance with IRB and institutional guidelines as above. Following transfer in 2% heat-inactivated FBS RPMI, tissue was cut into 1mm x 1mm cubes and placed in 0.25mg/ml liberase TH dissociation medium (Roche Diagnostics, Indianapolis, USA). Following further dissection, the tissue was incubated at 37°C for 1 hour at 600rpm. Samples were regularly triturated during the incubation period using a 1ml pipette, after which the digestion was stopped by the addition of 10% heat-inactivated FBS RPMI. The addition of ACK lysing buffer (ThermoFisher Scientific, Waltham, USA) following centrifugation at 500g for 5 minutes at room temperature, was performed twice in light of the lack of perfusion prior to nephrectomy. After centrifugation, the cell pellet was incubated with Accumax at 37°C for 3 minutes (Innovative Cell Technologies Inc, San Diego, USA), with 10% FBS RPMI again used for its subsequent neutralization. The resulting cell pellet was resuspended in 0.4% BSA/PBS and filtered using a 30um CellTrics filter (Sysmex America Inc, Lincolnshire, USA). Cell viability and concentration were determined using trypan blue, with 10,000 cells loaded into the 10x Genomics microfluidic system according to the manufacturer’s guidelines (10x Genomics, Pleasanton, USA).

### Droplet-based sn/scRNA-seq

Single nuclei or cells were partitioned into gel bead-in-emulsions (GEMs) and incubated to generate barcoded cDNA by reverse transcription. Barcoded cDNA was then amplified by PCR prior to library construction. Fragmentation, sample index and adaptor ligation, and PCR were used to generate libraries of paired-end constructs according to the manufacturer’s recommendations (10x Genomics, Pleasanton, USA). Libraries were pooled and sequenced using the Illumina HiSeq X system (San Diego, USA). Whenever feasible, we pooled 10x libraries on sequencing lanes to ensure that any individual sample was not confounded by batch (kidney section, day of sample collection, condition, timepoint) and were randomly distributed across lanes.

### Preprocessing of 10x droplet-based sequencing outputs

We used the *Cellranger* toolkit (v2.1.1, v3, 10X Genomics https://support.10xgenomics.com/single-cell-gene-expression/software/pipelines/latest/what-is-cell-ranger) to de-multiplex (*cellranger mkfastq*) the sequencing outputs, and for alignment (*cellranger count*) to the reference transcriptome (GRCh38 for human cells, GRCh38 pre-mRNA for human nuclei), and quantification of gene expression.

### Cell type identification and annotation

#### Quality control and normalization

All sc/snRNA-seq data analysis except co-embedding was done using the package *Seurat* (v2.3 (29) and v3.1 (30)). For the integrated analysis, we used the *merge* function to merge the expression count matrices. We used the *FilterCells* or *subset* function to retain cells that had read mapping to a minimum of 200 genes and raised the maximal mitochondrial threshold for cell inclusion to 40% mitochondrial gene reads as the kidney is a highly metabolically active organ. We normalized the data using total sum scaling followed by multiplication by a factor of 10^5^(TPX), and log-transformation using a pseudocount of 1 by the *NormalizeData* function to obtain log(TPX+1) values. Next, we identified high varying expression features using the *FindVariableGenes* function using default parameters for the biopsy cohort, and the *FindVariableFeatures* function with variance stabilization for the baseline integrated analysis. We computed the top 20 principal components (PCs) using the *RunPCA* function on the expression matrix composed of only the 2,000 most highly variable genes after mean centering and scaling using the *ScaleData* function.

#### Joint analyses of human cells from multiple donors

Batch effects are often detected by segregation of clusters by technical factors (*e.g*., donor origin, replicate, day of sequencing etc.) rather than expected biological identity. The single nuclei profile initially separated by donor. We used the *Harmony v1* (31) R package to co-embed human singlecell and single-nucleus RNA-seq data prior to clustering and visualization. Each donor was used as a separate batch.

#### Cell type identification

We performed clustering using the *FindClusters* function on the top 20 Harmony dimensions at a resolution of 1. *FindClusters* builds a shared *k*-nearest neighbor (*k*-NN, k=20) graph followed by community detection to determine clusters. We first used a resolution of 1, followed by merging of subsets in case of overclustering to retain major cell types after computing marker genes by differential expression (below). For 2-D visualization of the data by way of Uniform Manifold Approximation and Projection for Dimension Reduction (UMAP), we ran the *RunUMAP* function on the 20 Harmony dimensions, and with default parameters.

#### Cell type annotation and signatures

We used the *FindAllMarkers* function with default parameters to compute highly differentially expressed (DE) genes distinguishing each cluster from all other cells using the Wilcoxon Rank-Sum text with Benjamini-Hochberg FDR adjustment (32). We generated a database of literature-derived genes for kidney parenchymal, stromal and immune cell types. We annotated clusters by checking the presence of literature-derived, cell type-specific genes among the top DE genes when possible. Cells were annotated at the level of broad cell types.

### Modeling age and sex effects on *ACE2* expression

We first fit a mixed effect model (Eq 1) that explicitly models donor and dataset or center variability. Because of its complexity for the current dataset, it likely captures the most prominent effects. So we also fit a less complex fixed effects model (Eq 2) without modeling donor variability. A pseudo-bulk analysis at the donor level was used to validate the direction of effect sizes. For both models, datasets were held out to determine robustness of the model results to dataset variability.

To test the effect of age and sex on *ACE2* expression, we fit a Poisson mixed effects model to the integrated analysis dataset:

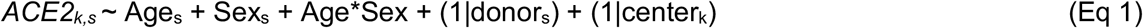

where *ACE2* is the gene expression of ACE2 in cell *k* and donor *s* in units of UMI counts. The total number of UMIs was added as an offset after scaling to have mean 1 and log-transformed. Age and Sex denote the age and sex of the donors in numerical years and binary “Female (reference)/Male”, respectively. The donor and dataset were modeled as nested random effects to account for batch effects. An interaction term was added to model the relationship between age and sex. We used the *glmer* function in the *lme4* (33) package to fit the model and compute p-values for individual coefficients.

The combined effect of sex and age can be summarized as: β_age_ + β_sex_ + sex*age*β_age:sex_ where β_age:sex_ is the interaction term and age is the numeric age, and sex is a binary variable (0 for females, 1 for male).

The sex effect can be summarized as: β_sex_ + age*β_age:sex_

The age effect can be summarized as: β_age_ + sex*β_age:sex_

For females (sex=0), the effect size is β_age_

For males (sex=1) the effect size is β_age_ + sex*β_age:sex_

*glmer* only computes uncertainty for and tests the individual coefficients. We computed uncertainties for the combined coefficients for age and sex effects by computing C^T^\***Σ**\*C where C is the vector of independent variables of interest, and **Σ** is the variance-covariance matrix of the fitted model. The combined effects and computed standard errors were subjected to a Wald test of statistical significance. Multiple hypothesis test correction was performed using the method of Benjamini-Hochberg (32) (FDR=5%). We performed leave one out cross validation (LOOCV) at the level of datasets to validate effect size direction and statistical significance.

To validate the direction of effects, pseudo-bulk analysis was performed by computing the mean *ACE2* expression count and mean UMI count across cells per cell type per donor. Using donors as observations, and the dataset as a random covariate, we fit the following Poisson mixed effect model to each cell type with the log-transformed mean UMI count as offset:

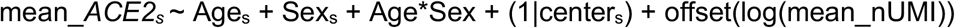

For capturing less complex effects, we fit the following Poisson regression model (5) with fixed effects and log-transformed UMI counts as offset:

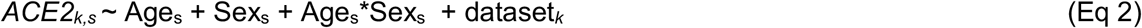

We used the *glm* function in R to fit the model. Pseudo-bulk analysis was performed by taking the mean counts as described above.

### Testing association between RAAS blockade and *ACE2* expression

To test the association between ACEi/ARB use and *ACE2* transcription, we fit the following Poisson mixed effects model to each cell type in the biopsy cohort:

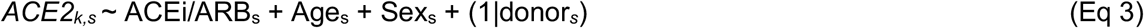

where *ACE2* is the gene expression of ACE2 in cell *k* and donor *s* in units of UMI counts. The total number of UMIs was added as an offset after scaling to have mean 1 and log-transformed. ACEi/ARB is the binary status “Yes/No” of whether the donor *s* was on the drug or not. Age and Sex denote the age and sex of the donor *s* in numerical years and binary “Female (reference)/Male”, respectively. The donor was modeled as a random covariate to account for batch effects. We used the *glmer* function in the *lme4* (33) package to fit the model and compute p-values. Multiple hypothesis test correction was performed using the method of Benjamini-Hochberg (32) (FDR=5%).

For the fixed effects model, we used the glm function in R to fit the following Poisson regression model:

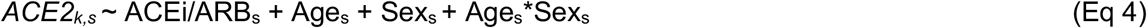

Pseudo-bulk analysis was performed as described above.

To model pathology, we fit the following two models:

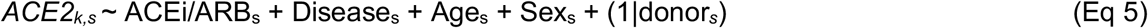

Because of its complexity, we could only fit (Eq 5) to 5 cell types.

Next, we fit a simpler model to determine disease-specific effects to all cell types with at least 5 *ACE2*^+^ nuclei.

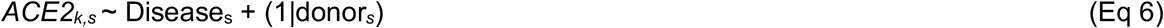

### Plotting and visualization

All analysis was performed in the R statistical computing environment (v3.6). We used the R visualization packages *ggplot2 v3.2.1* (34), *cowplot v0.9.4* (35), *ggpubr v0.2.5* (36) and *patchwork v1.0.0.9* (36, 37) for generation of boxplots, violin plots, proportional bar plots, UMAP visualization and dotplots. Data was wrangled using the *tidyverse* (38) framework (*dplyr v0.8.3, tidyr v1.0*).

### Availability of datasets and code

The “integrated analysis” baseline dataset (except the unpublished cohort) will be available as a resource on the Single-Cell Portal (SCP937)). Raw expression counts for *ACE2* for the unpublished nephrectomy and biopsy cohorts will also be available in the same study. Code used in the analysis will be made available at https://github.com/ayshwaryas/kidney_scmetaanalysis_c19

## Author contributions

K.A.V., M.S and J.W. performed experiments with assistance from D.D., M.C., K.K., J.P., L.N. and D.D. A.W. sampled nephrectomy tissue and performed histological analysis. A.S. and A.R. designed the analysis framework. A.S. performed the computational analysis with input from A.R. A.S., A.R., M.L and K.G. interpreted the statistical analysis. A.S., K.A.V., A.R. and A.G. wrote the manuscript. O.R-R., A.R. and A.G. supervised the project and all authors read and approved the manuscript.

## Acknowledgements

We thank Amanda Snook and the Division of Urology at Brigham and Women’s Hospital for their assistance with nephrectomy tissue sampling, and the patients involved in this study. We thank our colleagues Paul Hoover, Sanjay Jain, Arnon Arazi, William Apruzzese, Haojia Wu, Parker Wilson, Benjamin Humphreys, Rajasree Menon, Jeffrey Hodgin, Matthias Kretzler and Benjamin Stewart for their quick responses to questions on their published datasets. We gratefully acknowledge the commitment of the entire Kidney HCA network to data sharing and collaboration. This work was supported by the Chan Zuckerberg Initiative as part of the Human Cell Atlas Consortium.

## Supplementary materials for this manuscript include the following

### Dataset legends

Dataset S1: Table of datasets used in the integrated analysis with information on study center, data accession links, study citation, dataset characteristics (number of donors), and sc- or snRNA-seq technology.

Dataset S2: Results of integrated analysis to test the effects of age and sex on *ACE2* expression. Statistics of estimated fixed and random effects for the (1) mixed effect Poisson regression model, (2) Fixed effect Poisson regression model, (3) pseudo-bulk analysis and (4) cross validation are shown in separate sheets. Cross-validation results are shown for the significant effects ascertained from the full data.

Dataset S3: Results of association test between RAAS blocker use and *ACE2* expression. Statistics of estimated fixed and random effects for the (1) mixed effect Poisson regression model, (2) Fixed effect Poisson regression model and (3) pseudo-bulk analysis are shown in separate sheets.

**Supplementary Figure 1:**
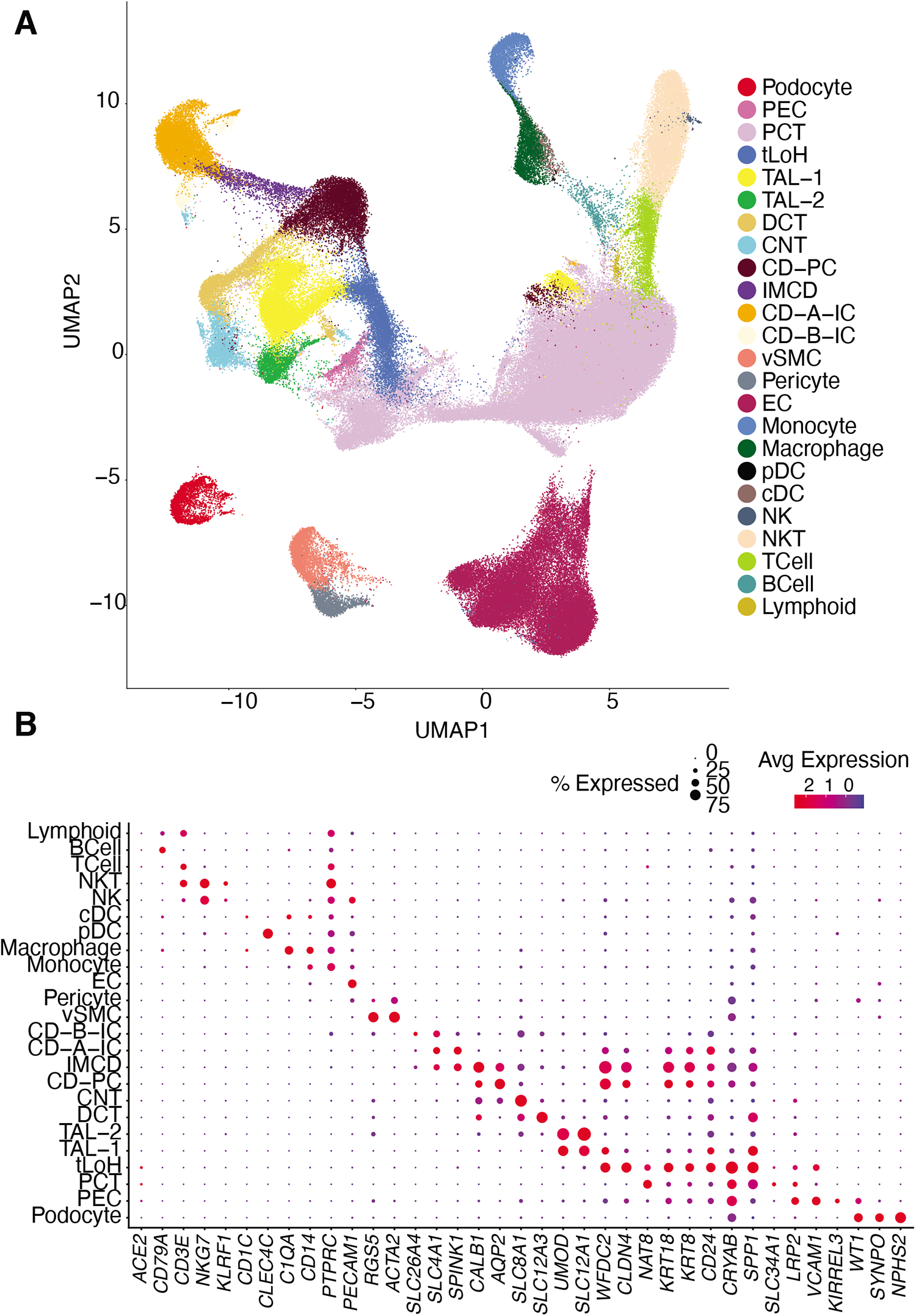
Integrated analysis dataset spans 24 broad kidney cell subsets. (**A**) Co-embedding. UMAP of cell profiles (dots) colored by annotated cell subset defined by clustering (**Methods**). (**B**) Cell subset annotations. Mean expression (log(TPX+1), dot color) of each marker (columns) in expressing cells and proportion of expressing cells (dot size) across cell types (rows).

**Supplementary Figure 2:**
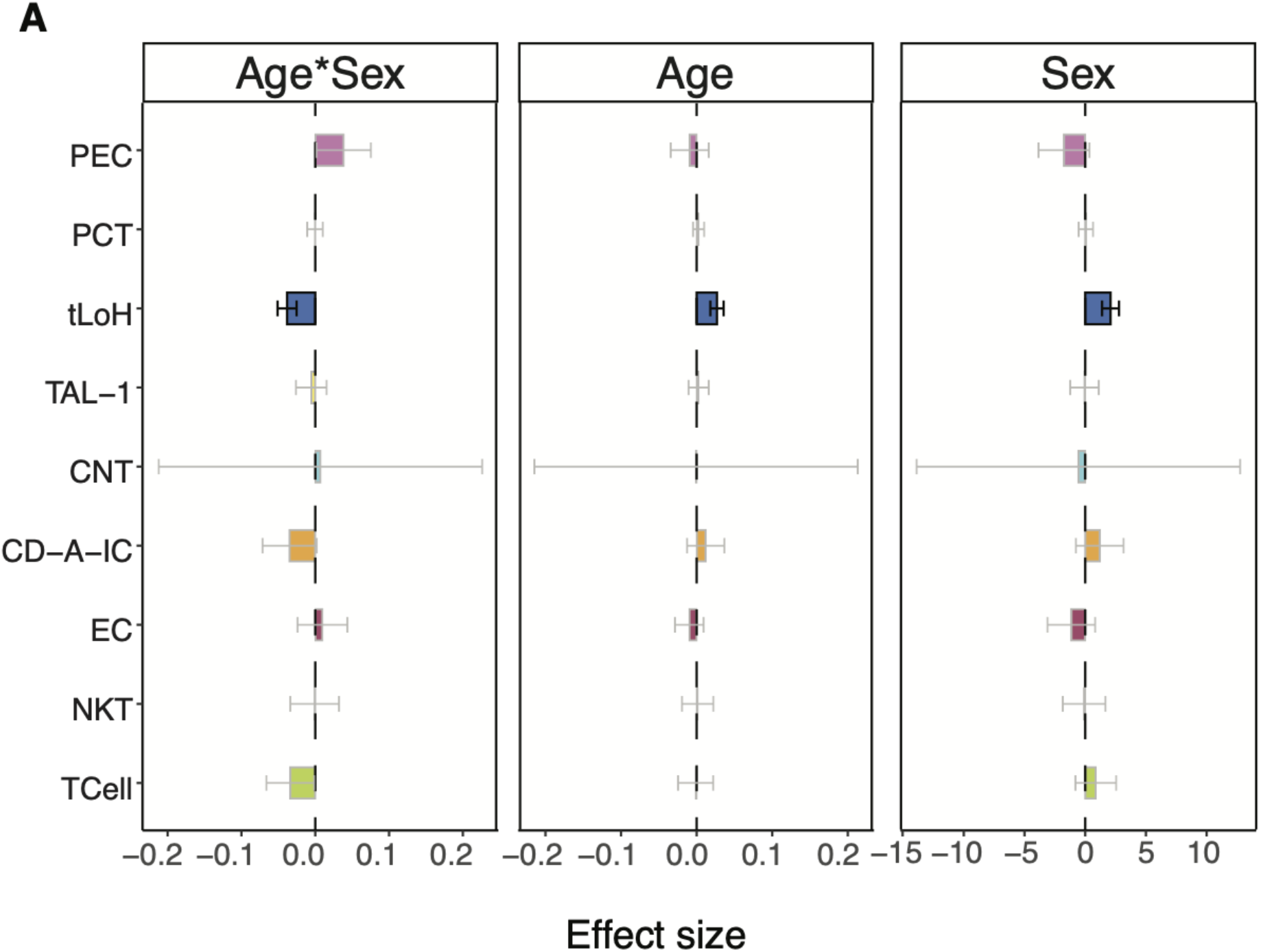
Association of ACE2 with age, sex and their interaction. The x-axis shows the effect size of the association in log-fold change (sex), or slope of log expression with age. Error bars represent standard errors around coefficient estimates. Statistically significant associations (FDR 5%) are outlined in black, others are in gray and colors are shaded lighter.

**Supplementary Figure 3:**
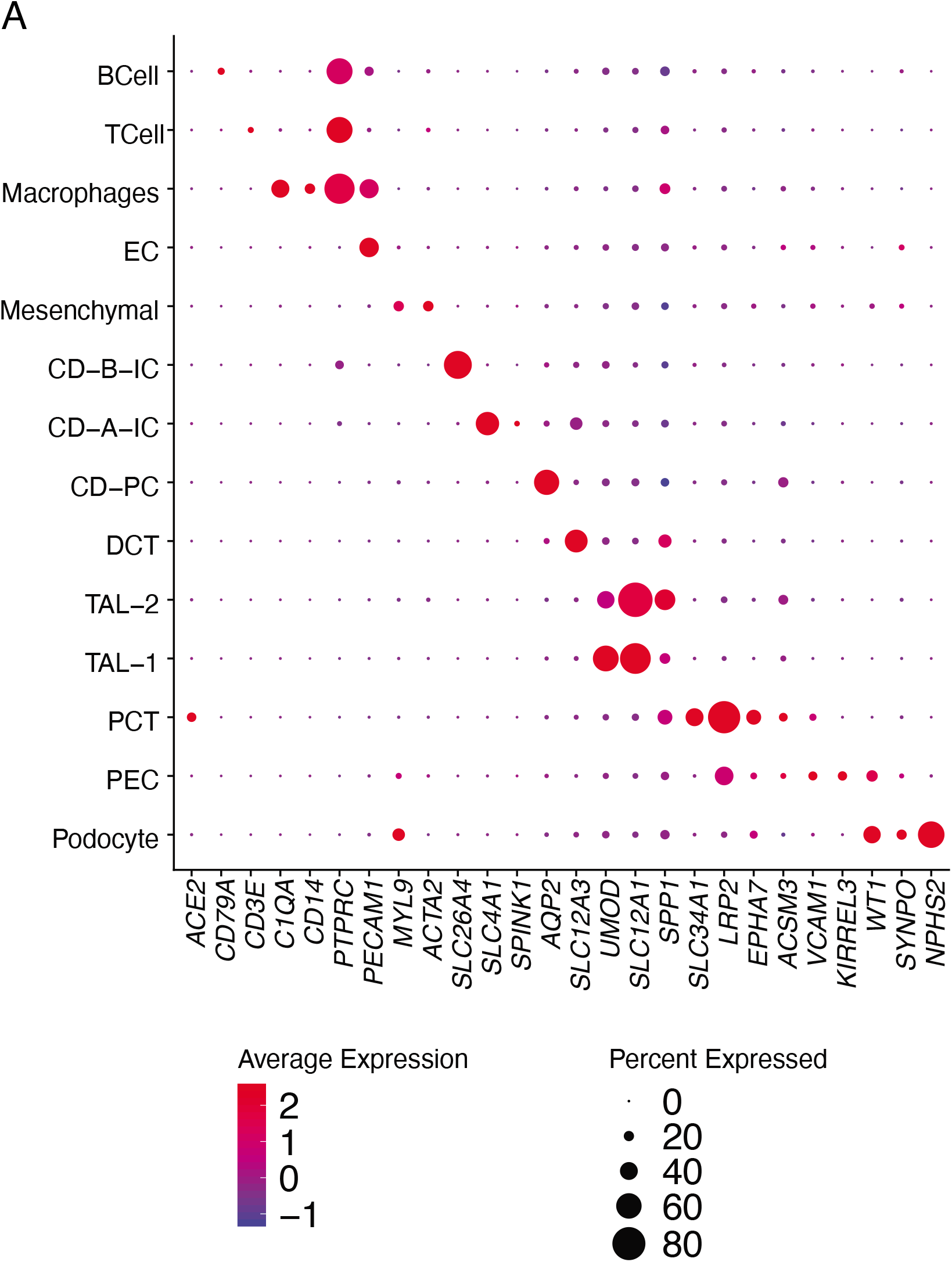
Cell subsets in the biopsy cohort. Mean expression (log(TPX+1), dot color) of each marker (columns) in expressing cells and proportion of expressing cells (dot size) across cell types (rows).

### Supplementary text

Human Cell Atlas Lung Biological Network - author list and affiliations

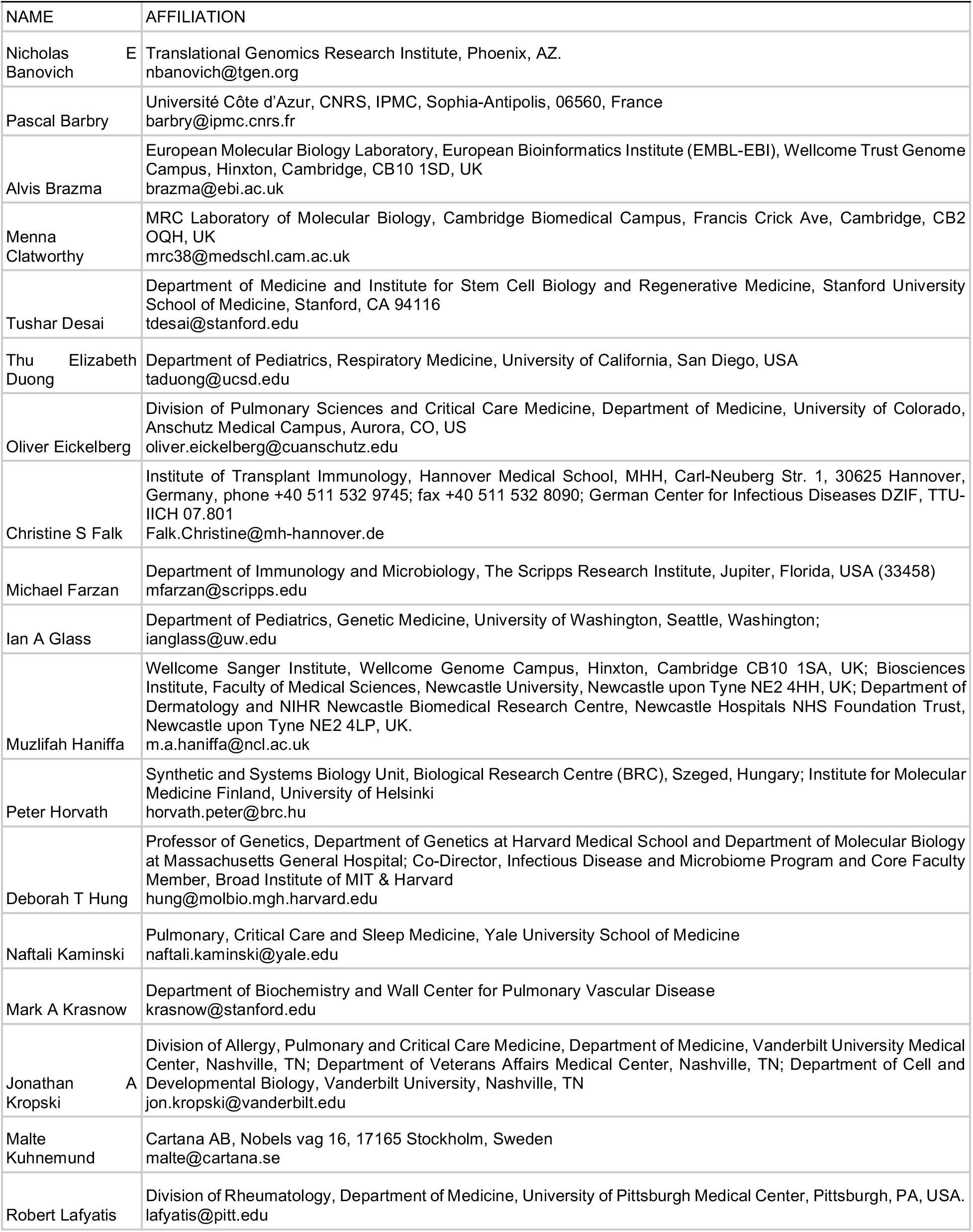
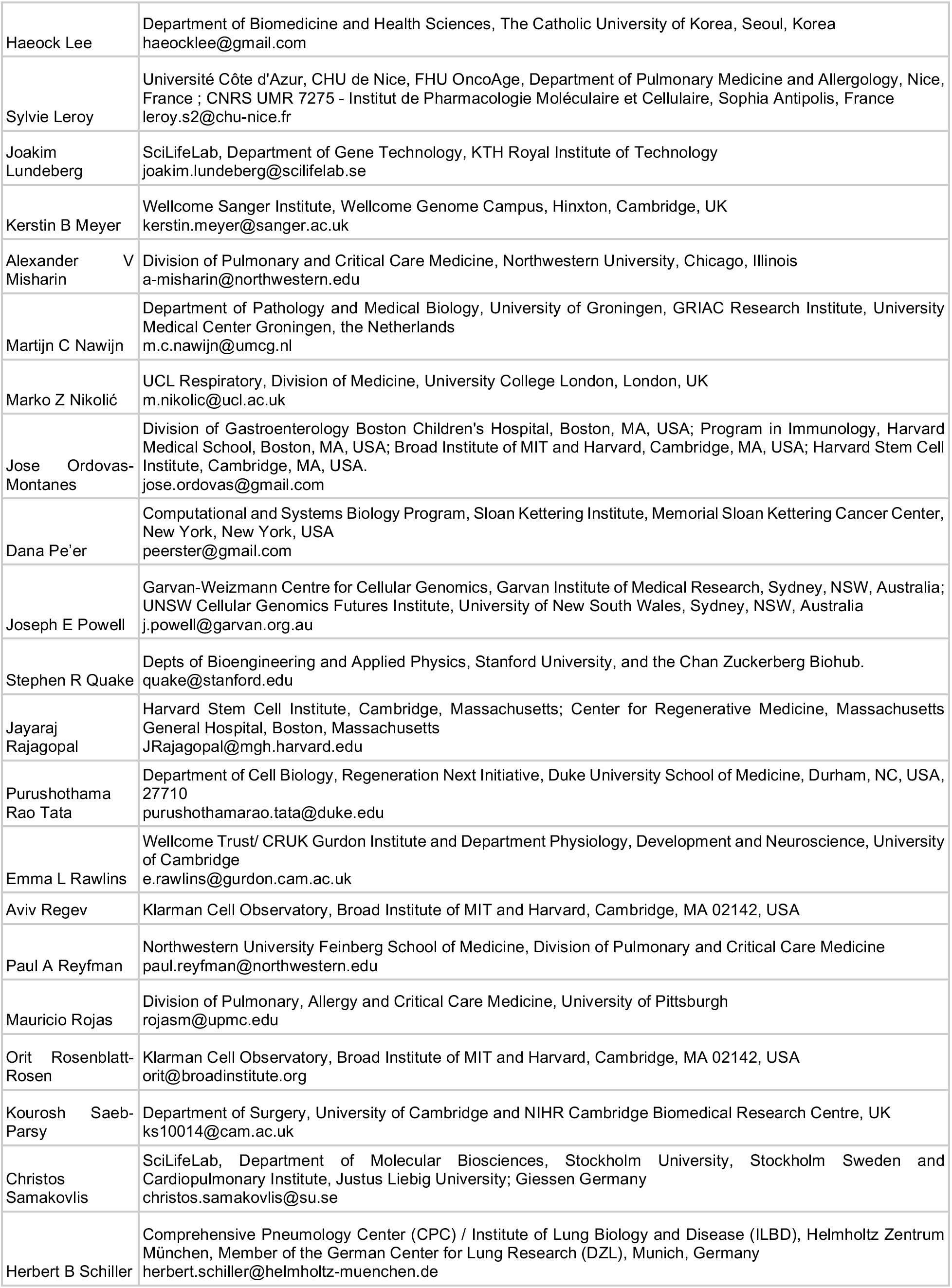
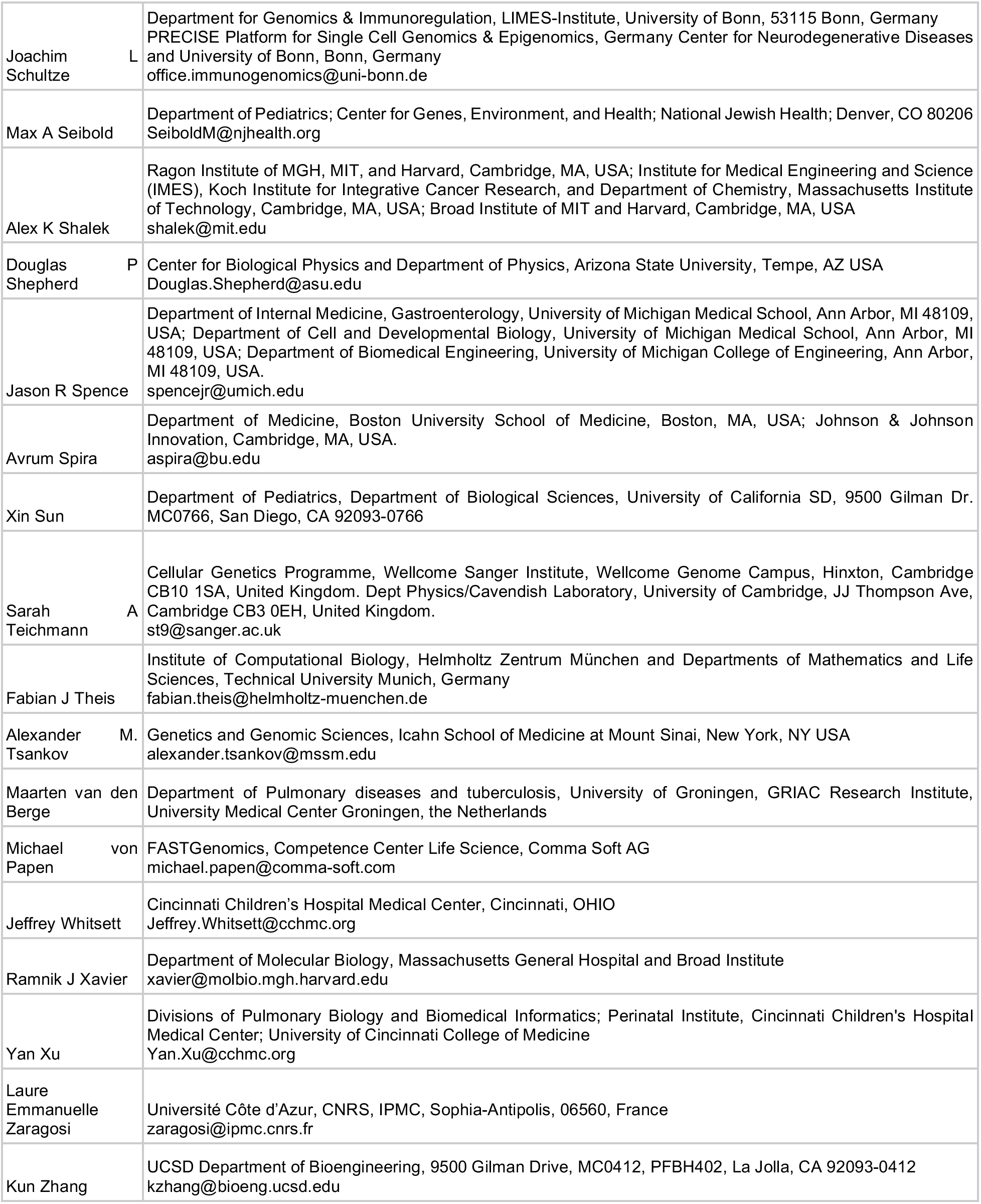

## References

1. W. Tai, et al., Characterization of the receptor-binding domain (RBD) of 2019 novel coronavirus: implication for development of RBD protein as a viral attachment inhibitor and vaccine. Cell. Mol. Immunol. (2020) https://doi.org/10.1038/s41423-020-0400-4.

2. J. Shang, et al., Structural basis of receptor recognition by SARS-CoV-2. Nature (2020) https://doi.org/10.1038/s41586-020-2179-y.

3. M. Hoffmann, et al., SARS-CoV-2 Cell Entry Depends on ACE2 and TMPRSS2 and Is Blocked by a Clinically Proven Protease Inhibitor. Cell 181, 271–280.e8 (2020).

4. D. Harmer, M. Gilbert, R. Borman, K. L. Clark, Quantitative mRNA expression profiling of ACE 2, a novel homologue of angiotensin converting enzyme. FEBS Lett. 532, 107–110 (2002).

5. The NHLBI LungMAP Consortium, The Human Cell Atlas Lung Biological Network, Integrated analyses of single-cell atlases reveal age, gender, and smoking status associations with cell type-specific expression of mediators of SARS-CoV-2 viral entry and highlights inflammatory programs in putative target cells (2020) https://doi.org/10.1101/2020.04.19.049254.

6. M. C. White, R. Fleeman, A. C. Arnold, Sex differences in the metabolic effects of the renin-angiotensin system. Biol. Sex Differ. 10, 31 (2019).

7. L. A. Ramirez, J. C. Sullivan, Sex Differences in Hypertension: Where We Have Been and Where We Are Going. Am. J. Hypertens. 31, 1247–1254 (2018).

8. K. Komukai, S. Mochizuki, M. Yoshimura, Gender and the renin-angiotensin-aldosterone system. Fundam. Clin. Pharmacol. 24, 687–698 (2010).

9. D. C. Villela, D. G. Passos-Silva, R. A. S. Santos, Alamandine: a new member of the angiotensin family. Curr. Opin. Nephrol. Hypertens. 23, 130–134 (2014).

10. S. Mizuiri, Y. Ohashi, ACE and ACE2 in kidney disease. World J Nephrol 4, 74–82 (2015).

11. E. Murray, M. Tomaszewski, T. J. Guzik, Binding of SARS-CoV-2 and angiotensin-converting enzyme 2: clinical implications. Cardiovasc. Res. (2020) https://doi.org/10.1093/cvr/cvaa096.

12. J. A. Jessup, et al., Effect of angiotensin II blockade on a new congenic model of hypertension derived from transgenic Ren-2 rats. Am. J. Physiol. Heart Circ. Physiol. 291, H2166–72 (2006).

13. Y. Ishiyama, et al., Upregulation of angiotensin-converting enzyme 2 after myocardial infarction by blockade of angiotensin II receptors. Hypertension 43, 970–976 (2004).

14. C. M. Ferrario, et al., Effect of angiotensin-converting enzyme inhibition and angiotensin II receptor blockers on cardiac angiotensin-converting enzyme 2. Circulation 111, 2605–2610 (2005).

15. A. Soro-Paavonen, et al., Circulating ACE2 activity is increased in patients with type 1 diabetes and vascular complications. J. Hypertens. 30, 375–383 (2012).

16. W.-J. Guan, et al., Clinical Characteristics of Coronavirus Disease 2019 in China. N. Engl. J. Med. 382, 1708–1720 (2020).

17. A. B. Patel, A. Verma, COVID-19 and Angiotensin-Converting Enzyme Inhibitors and Angiotensin Receptor Blockers: What Is the Evidence? JAMA (2020) https://doi.org/10.1001/jama.2020.4812.

18. R. Sommerstein, M. M. Kochen, F. H. Messerli, C. Gräni, Coronavirus Disease 2019 (COVID-19): Do Angiotensin-Converting Enzyme Inhibitors/Angiotensin Receptor Blockers Have a Biphasic Effect? Journal of the American Heart Association 9 (2020).

19. A. M. South, L. Tomlinson, D. Edmonston, S. Hiremath, M. A. Sparks, Controversies of renin–angiotensin system inhibition during the COVID-19 pandemic. Nature Reviews Nephrology (2020) https://doi.org/10.1038/s41581-020-0279-4.

20. G. Mancia, F. Rea, M. Ludergnani, G. Apolone, G. Corrao, Renin–Angiotensin–Aldosterone System Blockers and the Risk of Covid-19. N. Engl. J. Med. (2020) https://doi.org/10.1056/NEJMoa2006923.

21. H. R. Reynolds, et al., Renin-Angiotensin-Aldosterone System Inhibitors and Risk of Covid-19. N. Engl. J. Med. (2020) https://doi.org/10.1056/NEJMoa2008975.

22. Y. Cheng, et al., Kidney disease is associated with in-hospital death of patients with COVID-19. Kidney Int. 97, 829–838 (2020).

23. H. Su, et al., Renal histopathological analysis of 26 postmortem findings of patients with COVID-19 in China. Kidney Int. (2020) https://doi.org/10.1016/j.kint.2020.04.003.

24. B. Diao, et al., Human Kidney is a Target for Novel Severe Acute Respiratory Syndrome Coronavirus 2 (SARS-CoV-2) Infection. Infectious Diseases (except HIV/AIDS) (2020) https://doi.org/10.1101/2020.03.04.20031120.

25. E. A. Farkash, A. M. Wilson, J. M. Jentzen, Ultrastructural Evidence for Direct Renal Infection with SARS-CoV-2. J. Am. Soc. Nephrol. (2020) https://doi.org/10.1681/ASN.2020040432.

26. V. G. Puelles, et al., Multiorgan and Renal Tropism of SARS-CoV-2. N. Engl. J. Med. (2020) https://doi.org/10.1056/NEJMc2011400.

27. H. Wu, Y. Kirita, E. L. Donnelly, B. D. Humphreys, Advantages of Single-Nucleus over Single-Cell RNA Sequencing of Adult Kidney: Rare Cell Types and Novel Cell States Revealed in Fibrosis. J. Am. Soc. Nephrol. 30, 23–32 (2019).

28. M. Slyper, et al., A single-cell and single-nucleus RNA-Seq toolbox for fresh and frozen human tumors. Nat. Med. 26, 792–802 (2020).

29. A. Butler, P. Hoffman, P. Smibert, E. Papalexi, R. Satija, Integrating single-cell transcriptomic data across different conditions, technologies, and species. Nat. Biotechnol. 36, 411–420 (2018).

30. T. Stuart, et al., Comprehensive Integration of Single-Cell Data. Cell 177, 1888–1902.e21 (2019).

31. I. Korsunsky, et al., Fast, sensitive and accurate integration of single-cell data with Harmony. Nat. Methods 16, 1289–1296 (2019).

32. Y. Benjamini, Y. Hochberg, On the Adaptive Control of the False Discovery Rate in Multiple Testing with Independent Statistics. Journal of Educational and Behavioral Statistics 25, 60 (2000).

33. D. Bates, M. Mächler, B. Bolker, S. Walker, Fitting Linear Mixed-Effects Models Usinglme4. Journal of Statistical Software 67 (2015).

34. H. Wickham, Getting Started with ggplot2. Use R!, 11–31 (2016).

35. C. O. Wilke, “cowplot: Streamlined Plot Theme and Plot Annotations for ‘ggplot2’” (CRAN, 2019).

36. A. Kassambara, “ggpubr: ‘ggplot2’ Based Publication Ready Plots” (CRAN, 2020).

37. T. L. Pedersen, “patchwork: The Composer of Plots” (CRAN, 2019).

38. H. Wickham, et al., Welcome to the Tidyverse. Journal of Open Source Software 4, 1686 (2019).

